# Mantpy: a framework for extracellular matrix analysis in spatial proteomics

**DOI:** 10.1101/2025.06.04.657781

**Authors:** Mohamed Ghafoor, James E. Parkinson, Thao Pham, Sokratia Georgaka, Michael J. Haley, Elliot Jokl, Karen Piper Hanley, Judith E. Allen, Tara E. Sutherland, Magnus Rattray

## Abstract

Spatial proteomics technologies now profile cells and the extracellular matrix (ECM) together *in situ*. Yet analysis tools remain cell-centric, despite the ECM playing an essential role in health and disease. Here we present Mantpy, a framework that represents the ECM, and its interface with cells, as spatial graphs. Mantpy builds ECM graphs directly from matrix markers and links them with cell graphs for joint cell–ECM analysis, supporting graph statistics, explainable graph deep learning and visualisation. Mantpy extends from a single ECM marker to multiplexed cell–ECM panels. It recovers layered tissue architecture in the intestine from collagen IV alone, quantifies infection-associated ECM remodelling in mouse liver, and relates cells to the surrounding matrix in mouse lung, where cellular context recovers synthetically corrupted matrix labels. Released with ECM-inclusive datasets and interoperating with the scverse ecosystem and pytorch geometric. Mantpy brings the matrix into spatial analysis alongside the cell.

## Introduction

The extracellular matrix (ECM) is a dynamic, non-cellular scaffold of proteins, glycoproteins and proteoglycans that provides the structural foundation of multicellular organisms [1]. At the cellular level, the ECM regulates adhesion, migration and differentiation, whilst at the tissue level, ECM composition and organisation help determine biomechanical properties [2]. Its remodelling is a recurrent feature of disease, from the altered stroma surrounding tumours and the fibrosis that leads to organ failure to the matrix scaffolds organising granulomas [3–7]. Understanding this remodelling requires knowing which matrix components are present, how they are organised in space, and which cells they surround.

Spatial proteomics technologies such as imaging mass cytometry (IMC) [8] and co-detection by indexing (CODEX) [9] now routinely capture more than 40 markers *in situ*, with a single panel profiling cellular and ECM components together. Yet analysis of these images has focused on the cells. Current ECM tools annotate the matrisome [10, 11], the inventory of ECM and ECM-associated proteins, in bulk and single-cell data [12]; infer cell–matrix communication from transcriptomics [13]; analyse the matrisome in spatial transcriptomics [14]; or cluster multiplexed images at the pixel level [15].

Cells, by contrast, are routinely analysed as cell graphs: networks of nodes joined by edges. Each node is a cell carrying features such as cell type or protein expression; each edge links neighbouring cells and can carry features of that relationship, such as the distance between them. Cell graphs are widely used in spatial single-cell analysis through scverse packages such as Squidpy [16, 17]. They also serve as inputs to explainable graph deep learning, which can learn disease states from the graph [18] and visualise the important nodes and edges [19, 20]. Two approaches have brought the matrix into graphs, but only partly. ECM-aware cell graphs attach matrix measurements to cell nodes, so the matrix has no graph of its own and its organisation cannot be analysed directly [21, 22]. Fibre-network models graph the matrix without cells and from a single marker, so they cannot analyse cell–matrix relationships or cluster multiple ECM markers [23]. Spatial domain tools such as BANKSY [24], UTAG [25], CellCharter [26] and GraphCompass [17] build neighbourhood graphs over cells or spots in scverse data structures. They can be applied to ECM patches once a tool has created them, but none tiles the matrix itself or joins cells and matrix in one graph. Together, these tools leave three problems unsolved in combination: representing the ECM and its interface with cells as an explicit graph; supporting graph deep learning on that representation; and interoperating with the existing spatial-omics ecosystem (Supplementary Table 1).

Here we present Mantpy, an open-source framework that addresses these problems, supporting spatial analysis of the ECM independently of cells or jointly with them (Fig. 1). At its core is the ECM graph, which represents spatial patches as nodes joined by edges and can be linked to a cell graph to form a joint cell–ECM graph, turning matrix organisation and cell–matrix association into quantities that can be computed and compared across conditions. Because spatial panels typically devote few channels to the matrix, the representation works with as little as one ECM marker and scales up to multiplexed cell–ECM panels. Mantpy stores patch measurements and one graph per relation in the same scverse data structures used by Scanpy and Squidpy [27–29], so ECM analysis operates on the same objects as single-cell and spatial analysis. It also exports graphs to PyTorch Geometric for graph deep learning, where models learn from tissue structure and can report which matrix features drive a prediction.

**Fig. 1.**
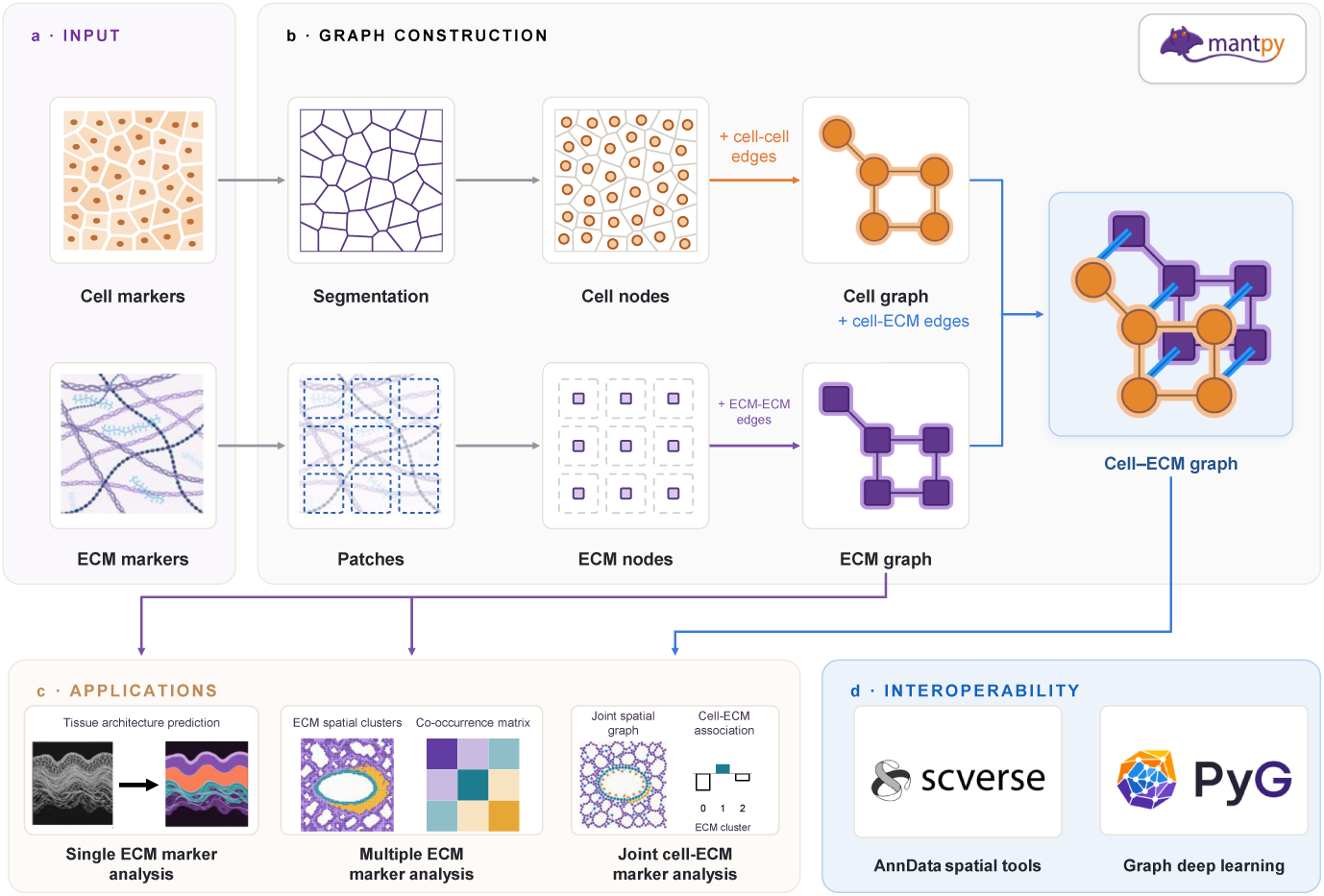
Overview of the Mantpy framework. **a**, Input. Mantpy takes one or more extracellular matrix (ECM) channels, alone or with cells for joint cell–ECM analysis. **b**, Graph construction. Segmented cells become cell nodes, and ECM channels are tiled into patches that become ECM nodes. Mantpy adds edges within each node type and then between them, giving a cell graph, an ECM graph and the joint cell–ECM graph. **c**, Applications. The ECM graph (purple arrows) and the joint cell–ECM graph (blue arrow) feed the analyses shown: single ECM marker analysis (tissue architecture prediction), multiple ECM marker analysis (ECM spatial clusters and their co-occurrence) and joint cell–ECM marker analysis (cell–ECM association). **d**, Interoperability. Mantpy exports graphs to PyTorch Geometric (PyG) for graph deep learning and stores them in scverse data structures (AnnData) for analysis alongside cellular workflows.

We demonstrate Mantpy on two imaging technologies, CODEX and IMC, with graphs of increasing complexity. In human small intestine, an ECM-only graph built from a single marker, collagen IV, recovers the layered architecture of the gut wall [30, 31]. In *Schistosoma*-infected mouse liver, multi-marker ECM graphs quantify infection-associated ECM remodelling and show where it occurs, and explainable graph deep learning highlights this compositional infection signal [7]. In healthy mouse lung, joint cell–ECM graphs built from ECM and cellular markers relate cell types to the surrounding matrix and recover synthetically corrupted ECM labels from cellular context [32]. Mantpy provides functions for constructing, visualising and analysing these graphs and is available at https://github.com/moeghaf/Mantpy; documentation, tutorials and ECM-inclusive datasets are available at https://mantpy.readthedocs.io/en/latest/ [33].

## Results

### Mantpy enables graph learning on the extracellular matrix

Mantpy constructs ECM graphs from single or multiple ECM markers. The graphs can be analysed on their own or jointly with a cell graph (Fig. 1a,b). It provides modules for ECM-inclusive dataset access, patch preprocessing, graph construction, analysis, graph learning and visualisation (Supplementary Table 2).

Unlike cells, the matrix is a continuous structure. Mantpy therefore adopts a patch-based unit of observation, an approach used widely in computational pathology and computer vision. The image channels containing matrix markers are divided into square spatial patches, and each patch becomes an ECM node carrying features derived from one or more channels. Features can include marker summaries and self-supervised embeddings. Foreground filters identify the patches to analyse; background patches can be removed or retained as unlabelled graph nodes, depending on the analysis. Neighbouring patches are joined by edges, forming the ECM graph. Clustering the patch features groups the patches into ECM clusters.

Cell graphs produced by upstream cellular segmentation and characterisation are retained, not replaced, so matrix analysis is added to an existing scverse workflow, alongside Scanpy [34] and Squidpy [16]. For joint analysis, the cell and ECM graphs are linked into a cell–ECM graph with two node types and three relationships: cell–cell edges encode adjacency among cells, ECM–ECM edges connect neighbouring patches, and cell–ECM edges link cells to nearby matrix (Fig. 1b). Each relationship is built independently, and both nodes and edges can carry features. Each graph feeds the applications in Fig. 1c: the ECM graph supports tissue architecture prediction from a single marker and ECM clustering with co-occurrence from several markers; the joint cell–ECM graph supports cell–ECM association. Mantpy stores graphs in AnnData and exports patch tables with their spatial coordinates and images to SpatialData [27–29]. Spatial analysis tools that accept AnnData use the standalone cell or patch tables. For graph deep learning, Mantpy exports graphs to PyTorch Geometric (PyG) [35] (Fig. 1d).

### Mantpy recovers tissue architecture from a single basement-membrane marker

In spatial proteomic analysis, tissue architecture is normally inferred from cellular composition: cells are usually segmented and characterised before spatial structure is assessed. We asked whether clustering an ECM graph built from a single marker could recover that structure without any cellular measurement. We used collagen IV, a basement-membrane protein present in every layer of the gut wall, in two human small-intestine CODEX images from two donors [9, 30, 31, 36].

Mantpy divided the collagen IV signal into ECM patches and used a self-supervised contrastive convolutional neural network (CNN) to learn a feature vector for each patch. These patch features became the node features of an ECM graph that connected each patch to its nearest neighbours. A graph masked autoencoder (GraphMAE) [37] learned node embeddings on this graph, and K-means grouped them into domains. Patch size and the graph, CNN and GraphMAE settings were tuned together for each image without anatomical labels, choosing the configuration whose clusters were most stable across training seeds (Supplementary Table 3). Manually annotated anatomical layers, five in image 1 and four in image 2, were used only for evaluation and to set the number of clusters (*K* = 5 and *K* = 4); choosing *K* by seed stability instead gave *K* = 2 in image 1 and *K* = 4 in image 2 (Supplementary Fig. 1a).

The methods compared differ in how much spatial context they use. Clustering pixel intensities uses none, clustering the CNN patch features uses the patch alone, and GraphMAE uses each patch together with its neighbours. Agreement with the manual annotation, measured by the adjusted Rand index (ARI) [38], increased along this axis: in image 1, ARI was 0.20 for pixel K-means, 0.45 for K-means on the CNN features and 0.90 for GraphMAE on the same features (Fig. 2a; Supplementary Fig. 1d). Image 2, which contains no muscularis propria, was harder for every method (0.01, 0.02 and 0.42; Fig. 2b).

**Fig. 2.**
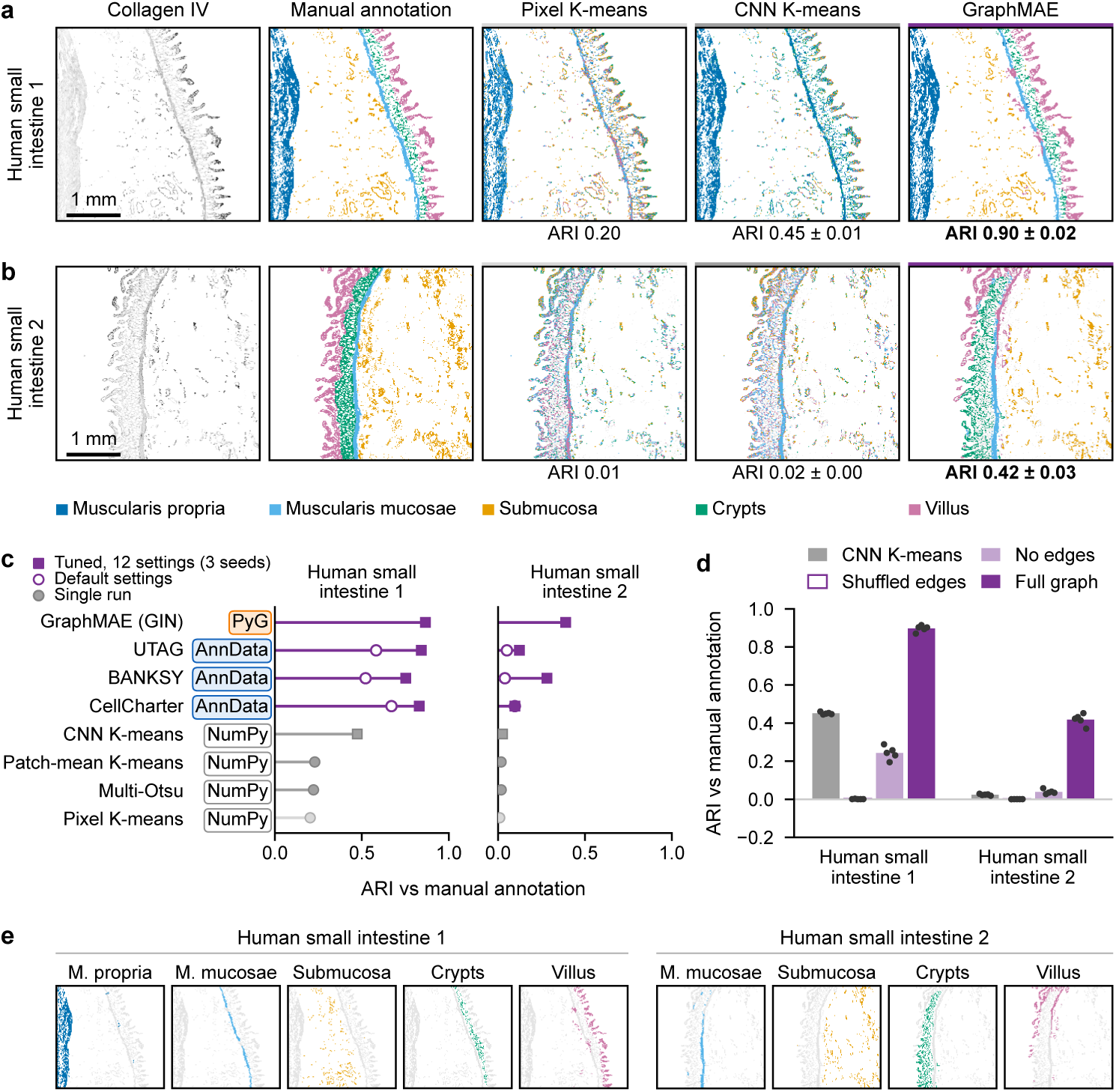
Collagen IV graphs recover tissue architecture in two human small-intestine images. **a**,**b**, Human small intestine 1 (**a**, *K* = 5) and 2 (**b**, *K* = 4): collagen IV signal, manual annotation and partitions by pixel K-means, CNN K-means on patch features and GraphMAE on the ECM graph. Values below the maps, ARI against the annotation (CNN K-means and GraphMAE, mean *±* s.d. over five training seeds; maps show the seed closest to the median). **c**, ARI of every method. Squares, tuned: the CNN was chosen by K-means seed stability and each spatial method chose among twelve neighbourhood settings without labels (means over three seeds; whiskers, s.d.); open circles, BANKSY, UTAG and CellCharter at default settings on the CNN of **a**,**b** (single runs); filled circles, single runs of the intensity baselines, which have no settings to tune. Tags, input passed to each method: NumPy array, Mantpy AnnData or PyTorch Geometric (PyG) graph. **d**, Graph controls: ARI of CNN K-means and of GraphMAE with shuffled edges, no edges or the full graph, with the CNN features and GraphMAE settings unchanged; bars, means over five training seeds; points, seeds. **e**, Each GraphMAE domain of **a** and **b**, shown against the other analysed patches (grey). Domains are named and coloured by their one-to-one match to the annotation; colours are the same in both images. Bars above the maps and in **c**: light grey, pixel; grey, patch; purple, graph methods. Scale bars, 1 mm. ARI, adjusted Rand index; CNN, convolutional neural network; GIN, graph isomorphism network; GraphMAE, graph masked autoencoder; *K*, number of clusters; M., muscularis.

Mantpy’s interoperability allowed the same patches and CNN features to be analysed in two ecosystems (Fig. 2c): as AnnData [28] by the scverse spatial tools BANKSY [24], UTAG [25] and CellCharter [26], and as an ECM graph exported to PyTorch Geometric (PyG) [35] by GraphMAE. Here the CNN and each method’s neighbourhood settings were chosen by the same seed-stability rule from twelve candidates each (Supplementary Fig. 1b,c), and the scverse tools were also run at their default settings. The comparison asks whether spatial context helps, not which spatial method is best. In image 1, every spatial graph method (ARI 0.52–0.87) agreed better with the annotation than every patch or pixel method (0.20–0.47). In image 2, ARI without a graph was at most 0.03, and GraphMAE (0.39) and tuned BANKSY (0.28) clearly exceeded it.

The spatial graph itself contributes this gain. With the CNN features, GraphMAE settings and final clustering held fixed, removing the edges lowered ARI from 0.90 to 0.24 in image 1 and from 0.42 to 0.04 in image 2, and shuffling them lowered it to 0.00 in both (Fig. 2d). The GraphMAE domains matched the annotated muscularis propria, muscularis mucosae, submucosa, crypts and villus (Fig. 2e). Because these layers extend well beyond a single patch, they favour representations that include neighbouring patches.

Mantpy therefore provides the patch representation, graph construction, graph deep learning and interoperability with scverse spatial tools and PyTorch Geometric needed to resolve tissue architecture from a single ECM marker.

### Mantpy describes ECM composition and organisation in infected mouse liver

A single marker recovers layered architecture, but matrix remodelling in disease is compositional as well as spatial: markers change individually and together, and the resulting groupings are distributed unevenly across a tissue. We used liver tissue from mice infected with *Schistosoma mansoni*, in which parasite eggs lodge in the liver and become encased in a granuloma. SOX9 is a transcription factor that promotes this fibrotic process [7]. We analysed six ECM markers in 24 mouse-liver IMC ROIs from eight mice of that study, crossing infection status with *Sox9* wild-type (WT) or knockout (KO) genotype, with two mice per group [7]. No cellular measurements were used.

Mantpy first grouped the ECM patches by their six-marker content, without using group information. As for the intestine, a contrastive convolutional encoder embedded each 10×10-pixel signal patch, here from all six marker channels, and K-means grouped the embeddings. The patch size, encoder settings and number of clusters were chosen by a label-free search, which selected six clusters (ECM 0–5), the largest number that remained reproducible across random seeds (mean pairwise adjusted Rand index across five seeds, 0.81; Supplementary Fig. 2 and Supplementary Table 4). The clusters differed in marker profile and in abundance between groups (Fig. 3a,b; per-mouse compositions and all ROIs, Supplementary Fig. 2). One cluster, ECM 5, was associated with infection. It was essentially absent from naive tissue (0–0.02% of ECM patches per mouse) but made up 4.0% of ECM patches in infected-KO and 15.2% in infected-WT mice (means of two mice; Fig. 3a). It had the highest fibronectin and heparan-sulfate signal of the six clusters, with high collagen I (Fig. 3b), and in infected ROIs formed compact regions (Fig. 3d,e). We therefore describe ECM 5 as infection-associated ECM remodelling; its composition and compact form are those expected of granuloma fibrosis [7], although granulomas were not annotated.

**Fig. 3.**
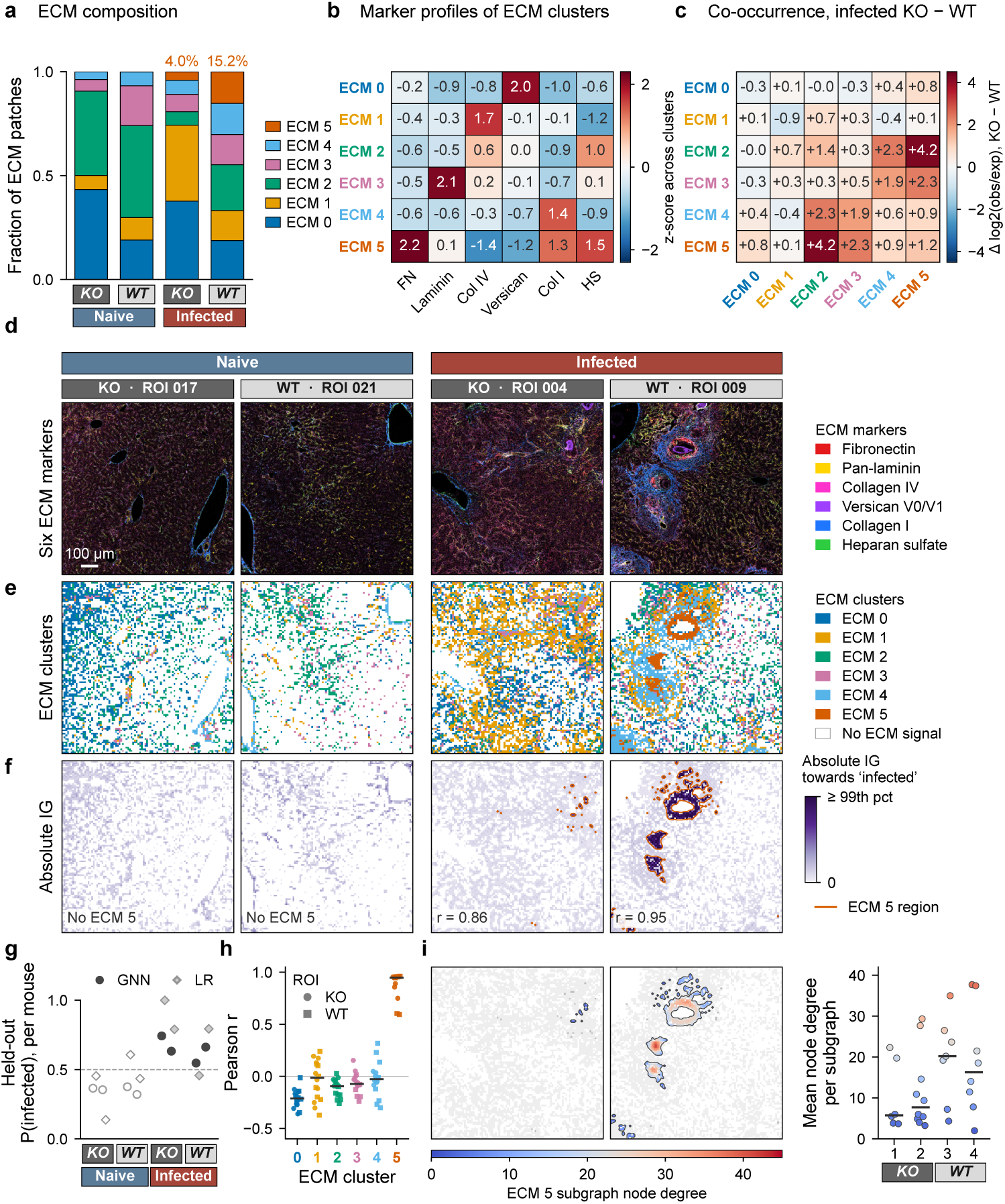
ECM composition and organisation in naive and *Schistosoma mansoni*-infected mouse liver. Imaging mass cytometry of six ECM markers in 24 regions of interest (ROIs) from eight mice, two per group (naive or infected; *Sox9* knockout (KO) or wild type (WT)). ECM patches of 10× 10 pixels were grouped into six clusters (ECM 0–5) without labels. **a**, Fraction of ECM patches in each cluster, mean of two mice per group; percentages, ECM 5. **b**, Mean marker intensity of each cluster, *z*-scored across clusters. **c**, KO − WT difference in cluster co-occurrence: log_2_(observed/expected) neighbour counts on the patch graph, expected from cluster abundance; red, higher in KO. **d**–**f**, One ROI per group (median number of ECM patches): six-marker composite (**d**), ECM clusters (**e**; white, no ECM signal) and absolute integrated-gradients (IG) importance towards infected (**f**; vermilion outline, ECM 5; *r*, Pearson correlation between *|*IG*|* and ECM 5 membership). **g**, Held-out P(infected) per mouse: graph neural network (GNN; circles) and logistic regression on cluster fractions (LR; diamonds); filled, infected; open, naive; dashed line, 0.5. **h**, Pearson *r* between *|*IG*|* and cluster membership in 16 infected ROIs (ECM 5: 14 ROIs with at least two ECM 5 patches); circles, KO; squares, WT; lines, medians. **i**, ECM 5 subgraphs (patches within 40 pixels joined; at least five patches) in the infected ROIs of **d**–**f**, coloured by node degree (grey, other ECM patches), and mean node degree per subgraph in each infected mouse (points, subgraphs; lines, medians). GNN values, mean of five networks trained without the held-out mouse, with the loss averaged within each ROI (**g**) or summed over patches (**f**,**h**). Scale bar, 100 *µ*m. Col I, collagen I; Col IV, collagen IV; ECM, extracellular matrix; FN, fibronectin; HS, heparan sulfate; P(infected), probability of infection; pct, percentile.

Because the ECM clusters are stored as node labels on the patch graph, how often they neighbour each other can be counted directly. We counted cluster pairs along the graph edges within each infected mouse and expressed each pair as log_2_(observed/expected), with the expected count set by the two clusters’ abundance. The largest KO-minus-WT difference, averaged over the two mice of each genotype, was between ECM 2 and ECM 5 (+4.2 log_2_ units): co-occurrence of these clusters was strongly depleted in both WT mice (−3.6 and −4.4) but close to expectation in both KO mice (+0.4 and 0.0; Fig. 3c and Supplementary Fig. 2). ECM 3 and ECM 5 followed the same pattern (+2.3). These values are descriptive: with two mice per genotype the smallest attainable two-sided exact *P* is 1*/*3, so no test is reported.

The same graphs support supervised learning, which we used to test whether a classifier and graph explainability tools would identify the infection-associated matrix without being told which cluster it is. A GraphSAGE classifier [39], receiving only ECM-cluster identity and graph connections, was trained with each mouse held out in turn on the frozen, cohort-defined partition (five-seed ensemble; loss averaged within each ROI). It assigned 8 of 8 held-out mice to the correct infection status (20 of 24 ROIs) and ranked every infected mouse above every naive mouse (ROC-AUC 1.00; Fig. 3g); tuning the classifier within each fold classified 7 of 8 (Supplementary Table 5). With two mice per group this result is limited: refitting the ensemble for every allocation of four infected labels gave *P* = 4*/*70 (*P* = 2*/*36 within genotype, the smallest attainable), and allocations labelling all KO or all WT mice as infected also reached ROC-AUC 1.00, so the representation separates genotype as completely as infection. Integrated gradients of the classifier trained with the loss summed over patches identified ECM 5 as its evidence: importance maps highlighted the ECM 5 regions, and in infected ROIs the correlation between node importance and ECM 5 membership had a median of 0.95 in KO and 0.96 in WT tissue, against medians between −0.23 and 0.00 for the other clusters (Fig. 3f,h). Cluster composition alone carried most of the signal, as a logistic regression on cluster fractions classified 6 of 8 mice (ROC-AUC 0.94; Fig. 3g). In this cohort the classifier therefore detects the presence of the remodelled matrix rather than its arrangement; the spatial differences between genotypes are described directly by the co-occurrence and subgraph analyses.

Graph statistics also describe how ECM 5 is organised. ECM 5 patches within 40 pixels were joined into connected subgraphs of at least five patches, and each subgraph was described by its size and mean node degree. Both KO mice had smaller and more fragmented ECM 5 subgraphs than both WT mice (median size 12.5 and 13.5 versus 61.0 and 68.5 patches; median mean node degree 5.8 and 7.7 versus 20.2 and 16.3; Fig. 3i). Mean degree is bounded by subgraph size, so these two descriptors are coupled. The genotypes had similar numbers of subgraphs (16 versus 15) although KO tissue contained 3.8-fold less ECM 5, so KO tissue had about four times as many subgraphs per ECM 5 patch. In the absence of *Sox9*, the remodelled matrix therefore formed smaller, more fragmented regions, consistent with SOX9’s role in granuloma fibrosis [7].

Together, the six-marker analysis separates the compositional and the spatial aspects of matrix remodelling. Compositionally, infection introduces a fibronectin- and heparan-sulfate-rich remodelled matrix that is essentially absent from naive liver and less abundant without *Sox9*. Spatially, that matrix is kept apart from the ECM 2 and ECM 3 clusters in WT but not KO mice, and it forms smaller, more fragmented regions in KO tissue. Each statement is read from the same patch graph: node labels for composition, edges for adjacency, connected ECM 5 subgraphs for organisation and node attributions for the classifier’s evidence. From a multiplexed matrix image, Mantpy therefore provides the preprocessing, graph analysis, deep learning and visualisation needed to quantify ECM remodelling and to show where it occurs in the tissue.

### Mantpy characterises cell–matrix associations from combined cell and ECM markers

The graphs so far used ECM markers alone. To relate cells to the matrix, Mantpy joins a cell graph (Fig. 4a) and an ECM graph (Fig. 4b) through cell–ECM edges into one cell–ECM graph (Fig. 4c), in which each cell is linked to the matrix patches around it. We built these graphs for six healthy mouse-lung IMC ROIs from two mice, with 12 annotated cell types and ten ECM markers [32]. Mantpy kept the existing cell annotations and clustered the ECM patches: the silhouette score selected four K-means components, one of which was background, leaving three signal clusters. Marker profiles and spatial locations supported annotating them as alveolar (ECM 0), bronchovascular-wall (ECM 1) and adventitial (ECM 2) matrix [40] (Fig. 4d). The cell, ECM and cell–ECM graphs of all six ROIs are shown in Supplementary Fig. 3.

**Fig. 4.**
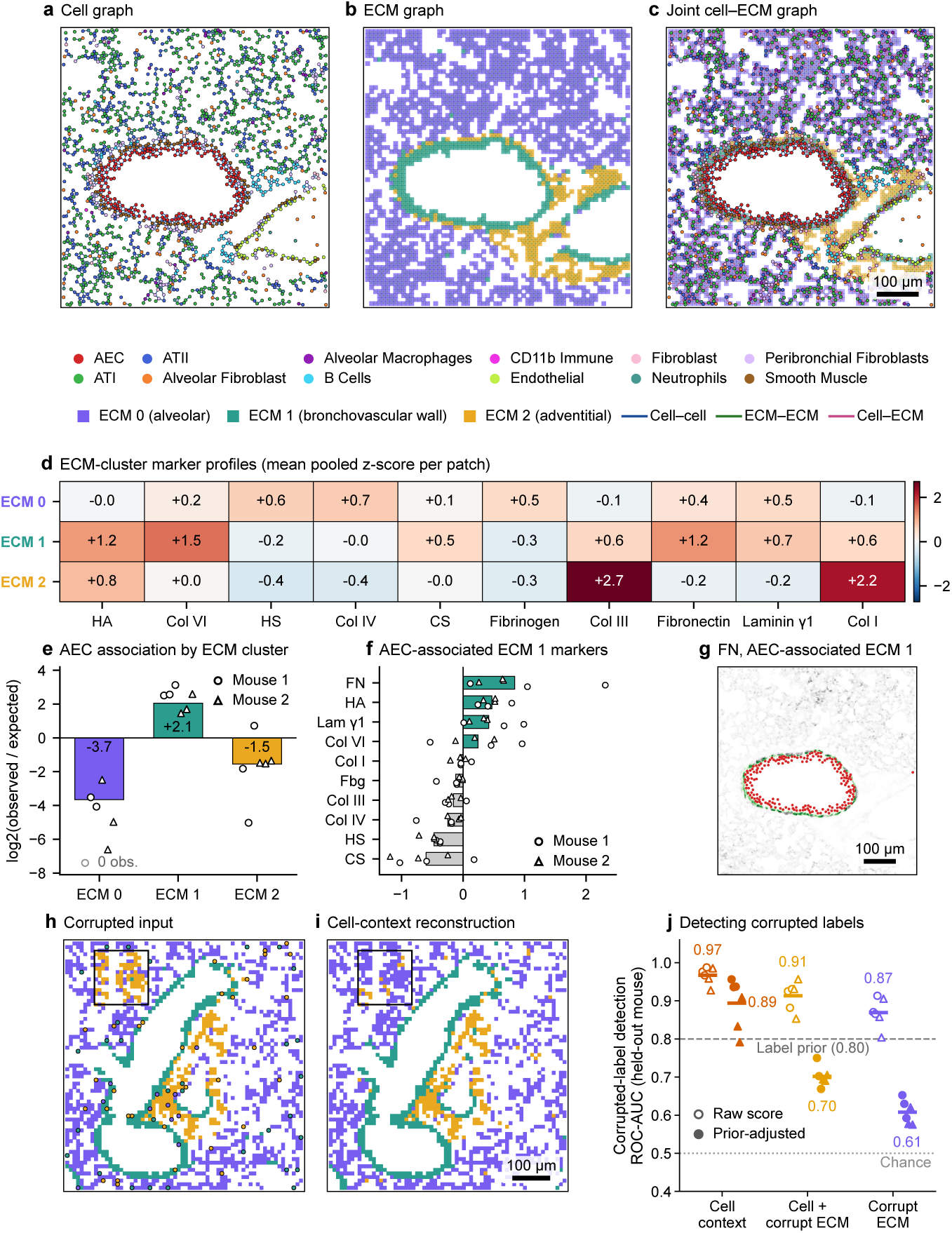
Cell–matrix association and recovery of ECM labels from cell context in healthy mouse lung. Imaging mass cytometry of six regions of interest (ROIs) from two mice, with 12 annotated cell types and ten ECM markers; 10 × 10-pixel ECM patches form three clusters (ECM 0, alveolar; ECM 1, bronchovascular wall; ECM 2, adventitial). **a–c**, Cell (**a**), ECM (**b**) and joint cell– ECM (**c**) graphs of one ROI. **d**, Mean *z*-scored marker intensity of the patches in each cluster; red, above and blue, below the mean. **e**, AEC association with each ECM cluster, log_2_(observed/expected) AEC–ECM edges pooled over ROIs (bars) and per ROI (circles, mouse 1; triangles, mouse 2; grey circle, 0 obs., ROI without an observed edge). **f**, Mean *z*-scored intensity of each marker in ECM 1 patches with minus without an AEC edge; bars, means over ROIs; symbols as in **e**. **g**, Fibronectin in AEC-associated ECM 1 patches (green) over the full fibronectin channel (grey), with AECs (red); displayed after a 3 × 3-pixel median filter. **h**,**i**, One ROI of a held-out mouse after replacing 10% of the ECM-cluster labels (**h**; black squares, replaced blocks; black-edged circles, isolated replacements) and after reconstruction from cell context by the model trained on the other mouse (**i**). **j**, ROC-AUC for detecting replaced labels from cell context, joint context (cell + corrupt ECM) or corrupted ECM context alone, with each mouse held out in turn, raw (open) and prior-adjusted (filled); points, ROIs of the held-out mouse (mean of five corruption realisations; symbols as in **e**); lines and numbers, means; dashed line, label-prior baseline; dotted line, chance. Scale bars, 100 *µ*m. AEC, airway epithelial cell; ATI and ATII, alveolar type I and II cells; Col, collagen; CS, chondroitin sulfate; Fbg, fibrinogen; FN, fibronectin; HA, hyaluronan; HS, heparan sulfate; Lam *γ*1, laminin-*γ*1; ROC-AUC, area under the receiver operating characteristic curve.

Counting cell–ECM edges then measures which matrix each cell type contacts. Airway epithelial cells (AECs) were linked to the bronchovascular-wall cluster, ECM 1, more often than expected (log_2_ observed/expected +2.1) and to ECM 0 and ECM 2 less often (−3.7 and −1.5; Fig. 4e). Per ROI, the ratio was positive for ECM 1 in all six ROIs, negative for ECM 0 in all six (one ROI had no AEC–ECM 0 edge) and negative for ECM 2 in five of six. With six fields from two mice, these associations are descriptive; the associations of every cell type and their sensitivity to the cell–ECM edge rule are shown in Supplementary Fig. 3.

The same edges select the matrix that a cell type touches. Within ECM 1, patches with an AEC edge had higher fibronectin, hyaluronan (HA) and laminin-*γ*1 than patches without one (mean differences +0.84, +0.47 and +0.42 standardised units) and lower heparan sulfate, collagen IV and fibrinogen, in all six ROIs (Fig. 4f). Mapped back onto the image, these patches outline the airway epithelium (Fig. 4g).

Because cell–ECM edges link each patch to the cells around it, the cell types near a patch can also predict its ECM cluster, which we used to correct corrupted ECM labels. In a synthetic denoising task, 10% of the ECM-cluster labels in each ROI were replaced, half in spatially continuous blocks and half as isolated patches. A classifier then predicted each patch’s cluster from the labels of its graph neighbours alone: its connected cells (cell context), its neighbouring patches (ECM context) or both (joint context). Each mouse was held out in turn, so no ROI of the test animal was seen during training or selection; graph rules, features and classifier settings were chosen within the three ROIs of the training mouse (Supplementary Table 6). Across the six held-out ROIs, cell context detected the replaced labels with a mean ROC-AUC of 0.97 (range 0.93–0.99; Fig. 4j) and corrected 70% of them (range 52–78%), while mislabelling 2.1% of uncorrupted patches, which raised mean label accuracy from 0.90 to 0.95 (Fig. 4h,i shows one held-out ROI). Correction was less complete for the rarer bronchovascular-wall and adventitial clusters (48% and 33%) than for the alveolar cluster (80%). Holding out single ROIs instead, with the other two ROIs of the same mouse in training, gave a ROC-AUC of 0.98 and corrected 78% of replacements, and cleaning accuracy stayed at 0.965–0.970 when the foreground and cluster centres were also refitted within the training ROIs.

The matrix’s own neighbourhood was less informative. Joint and ECM context reached ROC-AUCs of 0.91 and 0.87, against 0.80 for a baseline that used only the training label frequencies, which is high because replaced labels are enriched for the rarer clusters (Fig. 4j). After adjusting for this prior, cell, joint and ECM context gave 0.89, 0.70 and 0.61, and cell context ranked first in every held-out ROI. Trained on uncorrupted ROIs, the ECM and joint models learn that neighbouring patches share a label, which misleads them inside corrupted blocks; cell context does not use ECM labels.

Together, the cell–ECM graph relates cells to the matrix in two ways. Its edges measure which matrix compartments a cell type contacts and how that matrix differs, here a fibronectin- and HA-rich bronchovascular matrix next to the airway epithelium, and they let the surrounding cells locate and correct corrupted ECM labels. From cell and ECM markers in one image, Mantpy therefore provides the joint graph, association measures, context models and visualisation needed to relate cells to the matrix around them.

## Discussion

In spatial proteomics, matrix markers are routinely measured but rarely analysed further. Mantpy addresses this by representing the matrix, and its interface with cells, as spatial graphs, which enables graph statistics and graph deep learning directly on the ECM, alone or jointly with cells. Mantpy is not specific to an imaging technology: we demonstrate it on CODEX and IMC, but any image of matrix markers, including conventional immunohistochemistry, could supply the input. Its contribution is the joint cell–ECM graph and the framework around it, from preprocessing and graph construction to analysis and visualisation, rather than any single downstream method.

The three use cases each illustrate a general point. First, tissue architecture can be recovered from the matrix alone. In human small intestine, an ECM graph built from collagen IV, a basement-membrane marker present throughout the tissue, recovered the layered anatomy of the gut wall without any cellular measurement, and clustering the same features with and without the graph, or with its edges shuffled, showed that the improvement came from the graph’s spatial structure. ECM analysis can therefore be performed even with a single marker. These images add a practical caveat: the number of domains was taken from the annotation, and without it seed stability chose a coarser two-domain partition in image 1, so in unannotated tissue a stability plateau should prompt inspection of the coarser and finer partitions rather than a single *K*.

Second, remodelling can be described by organisation as well as composition. In *Schistosoma*-infected liver, a fibronectin- and heparan-sulfate-rich ECM cluster with high collagen I, which we describe as infection-associated ECM remodelling, was present in infected tissue and essentially absent from naive tissue. In the two *Sox9*-KO mice this cluster was less abundant, formed smaller and more fragmented subgraphs, and bordered a second cluster about as often as expected from abundance, whereas the two clusters were strongly segregated in the two WT mice. These observations are consistent with the more diffuse liver injury reported in infected *Sox9* KO mice [7]. Such differences are graph statistics on the ECM: subgraph size and distribution measure fragmentation, cluster co-occurrence measures neighbourhood, and both can be computed directly once the matrix has a graph of its own. Explainable graph deep learning then mapped the evidence for the classifier’s predictions back onto the tissue, where it concentrated on the remodelled matrix cluster. A logistic regression on cluster fractions classified nearly as well, so here the classifier detected the cluster’s presence rather than its spatial arrangement.

Third, cells and matrix inform each other. Joint cell–ECM graphs in mouse lung made cell–matrix association a measurable quantity, with airway epithelial cells preferentially associated with bronchovascular-wall matrix, and carried information across the compartment boundary. Each cell type receives a profile of the matrix it associates with, and in the denoising task cellular context repaired deliberately corrupted matrix labels. Repair of naturally erroneous measurements, missing channels or labels in new images was not tested; the label-reconstruction routine in mt.tl is therefore a benchmark of cross-compartment information flow on synthetic corruption, not a validated denoising utility for real data.

These applications used small cohorts, two intestine images, two mice per liver group and two lung mice, so they illustrate what the framework can measure rather than establish biological conclusions. Spatial imaging is also increasingly three-dimensional, and analysing the matrix together with cells in volumetric data is an open challenge. ECM and cell–ECM graphs could be built in three dimensions, but this remains future work. Releasing the ECM-inclusive datasets used here, each loading with a single Mantpy call [33], enables the community to develop ECM-focused algorithms. As such datasets accumulate, a shared representation would let matrix organisation be compared across tissues, diseases and cohorts, a step towards matrix atlases: reference descriptions of ECM organisation in health and disease. The self-supervised features used here could be extended into foundation models for ECM embeddings, trained across datasets rather than within single studies. The matrix is central to how tissues develop, repair and fail. As spatial panels include more ECM markers and spatial proteomics reaches the breadth of spatial transcriptomics, representing the matrix and its interface with cells as graphs will extend the unit of spatial analysis beyond the cell, to the matrix that surrounds it.

## Methods

### Data structures and interoperability

Mantpy stores data in the scverse containers AnnData and SpatialData [28, 29]. Cells and ECM patches have separate feature spaces and are kept in separate AnnData tables: X holds a compartment’s features, obs its identifiers and annotations, obsm its coordinates and embeddings, and obsp its sparse adjacency matrices; multidimensional edge features are stored in uns with their edge indices. The joint cell–ECM graph is a table of *n* cell and *m* patch rows with a node type annotation and no feature columns, so features of different dimensions are not padded into a shared variable space and the per-type tables are retained. Each relation is an (*n* + *m*) × (*n* + *m*) sparse matrix in obsp, with the *n × m* cell–ECM matrix *C* in the cross-type blocks, 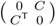. A SpatialData exporter stores each ROI image with its patch centres and links them to the patch table. Because the tables follow scverse conventions, Scanpy analyses run on the cell table [27, 34], an existing Squidpy cell graph can be reused [16], and BANKSY, UTAG and CellCharter receive the patch table.

### ECM patch representation and features

Mantpy lays a fixed grid of square, non-overlapping patches over the image, and each analysed patch becomes an ECM node. Patch size is a user-set parameter, chosen to match the imaging resolution and the scale of the structures of interest. Patch sizes and distances are given in image pixels, which are 1 *µ*m in the liver and lung IMC images and 0.37744 *µ*m in the intestine CODEX images. Foreground is selected by intensity or quantile thresholds on patch summaries or by masks of thresholded pixels, and after clustering `remove_background_patches` can assign the lowest-signal component to background; the dataset-specific filters are described below. Patches are described, per analysis, by a contrastive convolutional-neural-network embedding learned without labels [41] or by per-marker summary statistics for multiplexed panels. Clustering the features yields ECM clusters, stored as a node feature for graph learning. Clustering and model selection use scikit-learn [42]; image operations use scikit-image [43].

### Spatial graph construction

Mantpy represents a tissue section as one undirected, heterogeneous graph,

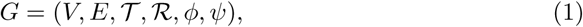

whose node set *V* = *V_C_ ⊔V_E_* contains cells *V_C_* = {*c*_1_, …, *c_n_*} with features 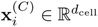 and ECM patches *V_E_* = {*e*_1_, …, *e_m_*} with features 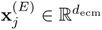, and whose edge set *E* = *E_CC_ ⊔E_EE_ ⊔E_CE_* contains cell–cell, ECM–ECM and cell–ECM edges. The maps *ϕ* : *V* → *T* = {Cell, ECM} and *ψ* : *E* → *R* = {*r_cc_, r_ee_, r_ce_*} assign node and relation types, so the cell, ECM and cell–ECM layers can be read from the same graph. Edges can also carry features.

Mantpy provides five rules for building relation-specific edges. Grid adjacency joins the four or eight neighbouring patches on the lattice and applies only to ECM nodes. Radius joins centres within the specified distance. *k*-nearest-neighbour graphs take the union of the queries from every node, with an optional distance cap; for cell–ECM edges the query runs from each cell to its *k* nearest patch centres, so a patch may receive many cell edges or none. Delaunay triangulation uses compartment centroids or, for cell–ECM edges, the pooled cell and patch centres with only cross-type edges retained, with optional jitter for ties on the regular lattice and a maximum edge length. The containing-patch rule links each cell to the patch it lies in and the eight surrounding patches that are present, so no cell is left without a matrix neighbour. A different rule may be set for each relation, and each graph is built through one call, with cell ecm joining the cell and ECM layers:

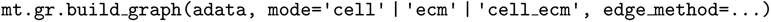

Background removal is a preprocessing step applied before or after graph construction, and an optional continuity check removes patch edges that cross background.

For an image of width *W* and height *H*, a complete square grid of patch side *s* contains approximately *WH/s*^2^ nodes. Grid, *k*NN (at most *kN* pairs) and Delaunay graphs have edge counts linear in the number of nodes *N*, whereas a large-radius graph can approach *N* ^2^ pairs. In a public-API scaling profile on repeated mosaics of one lung ROI, completed graphs reached 291,456 nodes and 3,454,380 directed edges; graph construction took up to 14.9 s and 20 GraphMAE updates 0.51–1.64 s on one GPU (Supplementary Fig. 4).

### Graph deep learning integration

Mantpy exports graphs as PyTorch Geometric `Data` or `HeteroData` objects [35], with edge indices, node features, positions and node identifiers in row order (and source identifiers where the graph builder recorded them); stored edge features are exported from patch graphs. Undirected relationships are represented in both directions, with separate cell-to-ECM and ECM-to-cell relations in `HeteroData`. Graph identifiers and optional fields are exported; other metadata remain in the source objects. The analyses used a GINE encoder [44], which uses Mantpy’s multidimensional edge features, for masked-feature autoencoding of ECM graphs, and SAGEConv [39], which supports homogeneous and bipartite message passing, for node classification. A GIN encoder without edge attributes was used in an ablation of the selected intestine configurations (identical graphs and CNN features, five seeds), in the intestine matched comparator control and in the scaling profile. Mantpy provides training modules for both analyses and remains compatible with other PyTorch Geometric models.

### Applications

The following subsections describe the dataset-specific preprocessing, graph construction and analysis for the three spatial proteomics applications.

#### Small-intestine collagen IV analysis

##### Data

We analysed the raw collagen IV channel of two HuBMAP CODEX small-intestine images from two donors, `HBM926.SHNZ.594` (image 1; proximal jejunum) and `HBM735.PSTF.274` (image 2) [9, 30, 31, 36]. Manually annotated tissue layers provided the evaluation reference: five in image 1 (muscularis propria, muscularis mucosae, submucosa, crypts and villus) and four in image 2, which contains no muscularis propria. Image 1’s manual square-tile labels were assigned to all pixels of their 32 × 32-pixel areas; image 2’s regions were annotated at native resolution and kept only where collagen IV intensity was nonzero. The known class counts fixed *K* = 5 and *K* = 4; otherwise the annotations were withheld from feature learning, graph construction, configuration selection and clustering, and used only for evaluation.

##### Patches and features

Non-zero intensities were clipped at the 99th percentile and rescaled to [0, 1]. A two-means threshold on patch means, fitted once at 32 × 32 pixels and then frozen, defined the foreground patches in both images, and both selected configurations used non-overlapping 32 × 32-pixel patches (13,376 patches in image 1 and 9,449 in image 2). A contrastive CNN trained with the SimCLR NT-Xent objective [41] learned a 96-dimensional embedding of each patch without labels.

##### Graphs and GraphMAE

Mantpy built an ECM graph with the patches as nodes, the standardised CNN embeddings as node features and edges to the 120 (image 1) or 160 (image 2) nearest neighbours, which carried standardised log-distance and orientation features. GraphMAE [37] used a four-layer, width-128 GINE encoder [44] with 50% masking, LayerNorm [45], a remasking decoder, jumping-knowledge aggregation [46] and a scaled-cosine reconstruction loss, and its embeddings were partitioned by K-means at the annotation’s *K*.

##### Configuration selection

The patch size, graph, CNN and GraphMAE settings of each image were chosen without the annotation by a multi-objective search (NSGA-II in Optuna) over 28 settings (Supplementary Table 3). Each configuration was fitted with three training seeds, each re-seeding the CNN, GraphMAE and K-means. Seed stability was the mean pairwise ARI on common 16-pixel foreground anchors, and coverage the fraction of the foreground inside analysed patches. Among configurations with coverage at least that of 32-pixel patches, the most stable was selected; configurations within 0.01 of it were refitted with two further seeds and compared at five seeds. The search evaluated 197 configurations for image 1 and 200 for image 2.

##### Comparison and scoring

We compared pixel K-means; Multi-Otsu thresholding [43, 47] and K-means of patch-mean intensities; K-means and Ward clustering of the CNN features; and BANKSY [24], CellCharter [26], UTAG [25] and GraphMAE as patch-graph methods. BANKSY, CellCharter and UTAG received the patch `AnnData` holding the CNN features and coordinates and built their own default neighbourhood graphs (BANKSY, pybanksy 1.3.5, and UTAG, commit 056cb87, reduced to 20 and 10 principal components and partitioned by K-means at the annotation’s *K*; CellCharter 0.3.7 fitting a Gaussian mixture with that *K*); GraphMAE received the ECM graph exported to PyTorch Geometric. In the matched comparison, the CNN was chosen among twelve architectures by CNN K-means seed stability, independently of GraphMAE, and each spatial method chose among twelve neighbourhood settings by the same stability rule before any reference scoring: BANKSY neighbourhood size (15, or once or twice the original neighbour count) with *λ* = 0.2, 0.4, 0.6 or 0.8; UTAG radius (20 *µ*m, median or 95th-percentile original edge length, or twice that percentile) with 10, 20 or 30 principal components; CellCharter graph (six neighbours, or half, once or twice the original count) with one to three aggregation hops; and GraphMAE, here with a GIN encoder, neighbour count (half, once or twice the original), depth (two or four layers) and masking (50% or 75%). Agreement was measured by ARI on the annotated pixels inside the analysed patches, identical for every method. Graph controls refitted GraphMAE with shuffled edges (every destination permuted) or no edges.

#### *Schistosoma*-infected mouse-liver analysis

##### Data and patches

The data are 24 IMC ROIs from eight mice of a published study [7], selected there on H&E-stained serial sections: naive or *S. mansoni*-infected, *Sox9* KO or WT, two mice per group and two ROIs per naive and four per infected mouse, with six ECM markers (fibronectin, pan-laminin, collagen IV, versican V0/V1, collagen I and heparan sulfate). Intensities were transformed with arcsinh (cofactor 1), scaled to each marker’s 99.5th percentile across ROIs and clipped to [0, 1]. Patches of 10 × 10 pixelswere described by each marker’s mean and standard deviation; after removing a low-signal background component (two-component K-means), 107,740 ECM signal patches remained.

##### Label-free ECM clusters

A contrastive convolutional encoder embedded the six-channel pixel block of each signal patch, and K-means clustered the standardised embeddings. The patch size, encoder settings and number of clusters *K* were chosen by a multi-objective search that maximised seed stability (mean pairwise adjusted Rand index between three seeds), coverage (fraction of the ECM signal covered at that patch size) and *K*. By a rule fixed before any encoder was trained, *K* was the largest value at which an eligible encoder reached a five-seed stability of at least 0.80, and the most stable encoder at that *K* was selected (10-pixel patches, *K* = 6, stability 0.81). No condition, genotype or mouse label was used, and the partition was frozen before downstream analysis (Supplementary Fig. 2 and Supplementary Table 4). Same-seed retraining of the encoder reproduced the partition with an adjusted Rand index of 0.96.

##### Infection-associated ECM remodelling cluster

ECM 5 was the cluster with the largest ratio of mean infected to mean naive per-mouse fraction (pseudocount 10*^−^*^3^) among clusters making up at least 1% of ECM patches in every infected mouse. This rule replaced the pre-specified largest-difference rule after the clusters were frozen, because that rule picked ECM 1, which is also common in naive tissue.

##### Graphs and co-occurrence

Each ROI’s graph joined ECM patches by the edges of the Delaunay triangulation of all patch centres. For each ROI, with symmetric adjacency *A* and one-hot cluster membership *L*, the ordered-endpoint counts are *O* = *L*^T^*AL*. Counts and node numbers *N_a_* were pooled within each mouse, and each cluster pair was expressed as log_2_(*O_ab_*/*E_ab_*) with *E_ab_* = *T p_a_p_b_*, where *p_a_* = *N_a_/*Σ*_b_ N_b_* is cluster abundance and *T* = Σ*_a,b_ O_ab_*. KO − WT differences are the mean of the two KO mice minus the mean of the two WT mice.

##### ECM 5 subgraphs

ECM 5 patches within 40 pixels were joined, and connected subgraphs of at least five patches were kept. Each subgraph was described by its patch count and mean node degree, which is bounded by *S −* 1 in a subgraph of *S* patches.

##### Classifier

A GraphSAGE classifier [39] received only each node’s one-hot cluster label and the graph edges and was trained with a class-weighted loss averaged within each ROI (two layers, 64 hidden units, mean aggregation, dropout 0.2; 120 epochs with Adam, learning rate 5 × 10*^−^*^3^, weight decay 10*^−^*^4^; settings not tuned on these data). Each mouse was held out in turn, with five seeds per fold, and a mouse’s probability of infection was the mean of its ROI probabilities (cut-off 0.5). Because the encoder and clusters were learned from all eight mice without labels, the cross-validation tests the classifier on this representation, not the full image-to-label pipeline. A logistic regression on ROI cluster fractions gave a composition-only reference. For tuning within each fold, 80 configurations of a 22-setting space (graph, layer and training settings; Optuna TPE) were scored by leaving out each training mouse in turn, and the best was refitted with five seeds (Supplementary Table 5). The permutation test refitted the ensemble for all 70 allocations of four infected labels. Integrated gradients [48] towards the infected class (zero baseline, 50 steps), from five classifiers trained with the loss summed over patches, were correlated with cluster membership in each held-out ROI.

#### Healthy mouse-lung cell–ECM analysis

##### Data and patches

We analysed six mouse-lung IMC ROIs from two mice, three per mouse [32]; the source study selected fields containing a small-to-medium airway, so the analyses describe airway-containing fields. Each ROI provided cell positions, labels for 12 cell types and images of ten ECM markers: hyaluronan, collagen VI, heparan sulfate, collagen IV, chondroitin sulfate, fibrinogen, collagen III, fibronectin, laminin-*γ*1 and collagen I. Images were divided into 10 × 10-pixel patches, each described by the mean of every ECM marker after standardising each marker over the pooled pixels of all ROIs. K-means with three to seven components was scored by the silhouette of the signal patches, which selected four; the lowest-signal component was removed as background, leaving 11,435 signal patches in three clusters, annotated as alveolar, bronchovascular wall and adventitial from their marker profiles and locations.

##### Graphs

The cell graph, built with Squidpy [16], connected all cell pairs within 15 pixels. The ECM graph connected adjacent patches on an eight-connected grid; background patches remained unlabelled nodes and were excluded from all counts. Cell–ECM edges joined each cell to its five nearest ECM patches within 15 pixels. The graphs for the denoising task were instead selected within each fold.

##### Cell–ECM association

For each cell type and ECM cluster, cell–ECM edges were counted in each ROI and compared with an expected count, the cell type’s total number of cell–ECM edges multiplied by the fraction of all cell–ECM edges incident on that cluster; observed and expected counts were summed over ROIs and expressed as log_2_(observed*/*expected). To profile the matrix associated with airway epithelial cells (AECs), the mean standardised intensity of ECM 1 patches without an AEC edge was subtracted from that of ECM 1 patches with one in each ROI, and the six ROIs were averaged with equal weight.

##### Synthetic corruption and reconstruction

A seeded generator replaced 10% of the signal-patch labels of each ROI, half within square blocks of side 50, 100 or 150 pixels and half as isolated patches, in five evaluation realisations. A multinomial logistic regression predicted each patch’s cluster from the cell-type composition of its connected cells (cell context), the cluster composition of its neighbouring patches (ECM context) or both, never from the patch’s own label; a label-prior baseline used only the training cluster frequencies, and a prior-adjusted detection score, −log[*P* (*y*_obs_ *|* context)*/P*_train_(*y*_obs_)], removed the effect of label frequency without changing cleaning. A patch was relabelled to the predicted cluster when its anomaly score, −log *P* (observed label *|* context), exceeded a threshold and the prediction differed from the observed label with a minimum confidence. Each mouse was held out in turn. Graph rules, features, regularisation and cleaning parameters were selected within the training mouse by nested Bayesian optimisation, with an inner loop holding out each of its three ROIs, maximising post-cleaning accuracy on separate tuning realisations (Optuna tree-structured Parzen estimator, 60 trials; selected configurations, Supplementary Table 6); the selected configuration was refitted on the training mouse’s pristine ROIs and scored once on each ROI of the held-out mouse. Scores were averaged over the five realisations within each ROI and then over ROIs with equal weight. Neighbouring marker intensities were excluded from the context comparison because they re-derive the cluster from the image. Controls held out single ROIs instead (150 trials, inner loop over the five training ROIs) and, with single ROIs held out, refitted the foreground and clusters within every training set.

#### Statistics

##### Intestine

Agreement with the manual annotation (ARI) is descriptive: GraphMAE and CNN K-means are reported as mean *±* s.d. over five training seeds, every method in the matched comparison over three seeds, and the other methods from one seed; no hypothesis tests were performed.

##### Liver

Co-occurrence and subgraph comparisons are descriptive: with two mice per genotype, the smallest attainable two-sided exact *P* is 1*/*3. The complete classifier protocol was refitted for every allocation of four infected and four naive labels; *P* is the fraction of allocations whose mouse-level ROC-AUC is at least the observed value, over all 70 allocations or the 36 that keep two infected and two naive mice per genotype.

##### Lung

Cell–ECM associations, ECM 1 marker differences and denoising results are descriptive: with six fields from two mice, effects are reported with their per-ROI consistency, and no hypothesis tests are reported.

## Supporting information

Combined supplementary

## Data availability

All data used in this study can be loaded through the Mantpy package. Human small-intestine CODEX datasets were generated by the HuBMAP Program and are available from the HuBMAP Data Portal under accessions HBM926.SHNZ.594 (image 1) and HBM735.PSTF.274 (image 2) [30, 31, 36]. Mouse lung and liver imaging mass cytometry datasets were obtained from Parkinson et al. [32] and Su et al. [7], respectively. Source data are provided with this paper.

## Code availability

Mantpy is available under the MIT licence at https://github.com/moeghaf/Mantpy and can be installed from PyPI. Documentation is available at https://mantpy.readthedocs.io. Notebooks and source code to reproduce the results are available at https://github.com/moeghaf/mantpy_reproducibility.

## Supplementary information

This paper is accompanied by Supplementary Information containing Supplementary Tables 1–6 and Supplementary Figs. 1–4.

## Author contributions

M.G. conceived the study with J.E.P., T.E.S. and M.R. M.G. and T.P. implemented Mantpy. J.E.P., T.E.S., E.J., K.P.H., S.G. and J.E.A. acquired the data. M.G. analysed the three datasets and wrote the paper. M.J.H. reviewed the paper. M.R. supervised the work. All authors read, corrected and approved the final paper.

## References

[1] Hynes, R. O. The evolution of metazoan extracellular matrix. Journal of Cell Biology 196, 671–679 (2012).

[2] Frantz, C., Stewart, K. M. & Weaver, V. M. The extracellular matrix at a glance. Journal of Cell Science 123, 4195–4200 (2010).

[3] Sorokin, L. The impact of the extracellular matrix on inflammation. Nature Reviews Immunology 10, 712–723 (2010).

[4] Sutherland, T. E., Dyer, D. P. & Allen, J. E. The extracellular matrix and the immune system: A mutually dependent relationship. Science 379, eabp8964 (2023).

[5] Henderson, N. C., Rieder, F. & Wynn, T. A. Fibrosis: from mechanisms to medicines. Nature 587, 555–566 (2020).

[6] Winkler, J., Abisoye-Ogunniyan, A., Metcalf, K. J. & Werb, Z. Concepts of extracellular matrix remodelling in tumour progression and metastasis. Nature Communications 11, 5120 (2020).

[7] Su, K. et al. Sox9 plays an essential role in myofibroblast driven hepatic granuloma integrity and parenchymal repair during schistosomiasis-induced liver damage. PLOS Pathogens 21, e1012928 (2025).

[8] Giesen, C. et al. Highly multiplexed imaging of tumor tissues with subcellular resolution by mass cytometry. Nature Methods 11, 417–422 (2014).

[9] Goltsev, Y. et al. Deep profiling of mouse splenic architecture with CODEX multiplexed imaging. Cell 174, 968–981.e15 (2018).

[10] Hynes, R. O. & Naba, A. Overview of the matrisome—an inventory of extra-cellular matrix constituents and functions. Cold Spring Harbor Perspectives in Biology 4, a004903 (2012).

[11] Naba, A. et al. The matrisome: in silico definition and in vivo characterization by proteomics of normal and tumor extracellular matrices. Molecular & Cellular Proteomics 11, M111.014647 (2012).

[12] Petrov, P. B., Considine, J. M., Izzi, V. & Naba, A. Matrisome AnalyzeR–a suite of tools to annotate and quantify ECM molecules in big datasets across organisms. Journal of Cell Science 136, jcs261255 (2023).

[13] Lamba, R., Paguntalan, A. M., Petrov, P. B., Naba, A. & Izzi, V. MatriCom, a single-cell RNA-sequencing data mining tool to infer cell–extracellular matrix interactions. Journal of Cell Science 138, jcs263927 (2025).

[14] Oshinjo, A., Chen, D., Petrov, P., Izzi, V. & Naba, A. MatriSpace: Identification and visualization of spatially resolved ECM gene expression patterns in health and disease. bioRxiv (2026). URL 10.64898/2026.04.26.720198. Preprint; not peer reviewed.

[15] Liu, C. C. et al. Robust phenotyping of highly multiplexed tissue imaging data using pixel-level clustering. Nature Communications 14, 4618 (2023).

[16] Palla, G. et al. Squidpy: a scalable framework for spatial omics analysis. Nature Methods 19, 171–178 (2022).

[17] Ali, M. et al. GraphCompass: spatial metrics for differential analyses of cell organization across conditions. Bioinformatics 40, i548–i557 (2024).

[18] Gunduz, C., Yener, B. & Gultekin, S. H. The cell graphs of cancer. Bioinformatics 20, i145–i151 (2004).

[19] Ying, Z., Bourgeois, D., You, J., Zitnik, M. & Leskovec, J. GNNExplainer: generating explanations for graph neural networks. Advances in neural information processing systems 32 (2019).

[20] Jaume, G. et al. Quantifying explainers of graph neural networks in computational pathology. Proceedings of the IEEE/CVF Conference on Computer Vision and Pattern Recognition (CVPR) 8106–8116 (2021).

[21] Bilgin, C. C., Lund, A. W., Can, A., Plopper, G. E. & Yener, B. Quantification of three-dimensional cell-mediated collagen remodeling using graph theory. PLOS ONE 5, e12783 (2010).

[22] Bilgin, C. C., Bullough, P., Plopper, G. E. & Yener, B. ECM-aware cell-graph mining for bone tissue modeling and classification. Data Mining and Knowledge Discovery 20, 416–438 (2010).

[23] Grapa, A.-I., Efthymiou, G., Van Obberghen-Schilling, E., Blanc-Féraud, L. & Descombes, X. A spatial statistical framework for the parametric study of fiber networks: Application to fibronectin deposition by normal and activated fibroblasts. Biological Imaging 3, e25 (2023).

[24] Singhal, V. et al. BANKSY unifies cell typing and tissue domain segmentation for scalable spatial omics data analysis. Nature Genetics 56, 431–441 (2024).

[25] Kim, J. et al. Unsupervised discovery of tissue architecture in multiplexed imaging. Nature Methods 19, 1653–1661 (2022).

[26] Varrone, M., Tavernari, D., Santamaria-Martínez, A., Walsh, L. A. & Ciriello, G. CellCharter reveals spatial cell niches associated with tissue remodeling and cell plasticity. Nature Genetics 56, 74–84 (2024).

[27] Virshup, I. et al. The scverse project provides a computational ecosystem for single-cell omics data analysis. Nature Biotechnology 41, 604–606 (2023).

[28] Virshup, I., Rybakov, S., Theis, F. J., Angerer, P. & Wolf, F. A. anndata: Access and store annotated data matrices. Journal of Open Source Software 9, 4371 (2024).

[29] Marconato, L. et al. Spatialdata: an open and universal data framework for spatial omics. Nature Methods 22, 58–62 (2025).

[30] Hickey, J., Caraccio, C. & Nolan, G. CODEX data from the small intestine of a 24-year-old white male (2023). URL 10.35079/HBM734.XBSR.357. Dataset.

[31] Hickey, J., Caraccio, C. & Nolan, G. CODEX data from the small intestine of a 30-year-old white male (2023). URL 10.35079/HBM363.CKHC.928. Dataset.

[32] Parkinson, J. E. et al. Extracellular matrix phenotyping by imaging mass cytometry defines distinct cellular matrix environments associated with allergic airway inflammation. Molecular Systems Biology (2026). URL 10.1038/s44320-026-00234-5.

[33] Ghafoor, M. Mantpy 0.2.0 tutorial datasets for spatial extracellular-matrix analysis. Zenodo (2026). URL https://zenodo.org/records/21538382. CC BY 4.0.

[34] Wolf, F. A., Angerer, P. & Theis, F. J. Scanpy: large-scale single-cell gene expression data analysis. Genome Biology 19, 15 (2018).

[35] Fey, M. & Lenssen, J. E. Fast graph representation learning with PyTorch Geometric. *arXiv preprint arXiv:1903.02428* (2019).

[36] HuBMAP Consortium. The human body at cellular resolution: the NIH Human BioMolecular Atlas Program. Nature 574, 187–192 (2019).

[37] Hou, Z. et al. GraphMAE: Self-supervised masked graph autoencoders. Proceedings of the 28th ACM SIGKDD Conference on Knowledge Discovery and Data Mining 594–604 (2022).

[38] Hubert, L. & Arabie, P. Comparing partitions. Journal of Classification 2, 193–218 (1985).

[39] Hamilton, W., Ying, Z. & Leskovec, J. Inductive representation learning on large graphs. Advances in neural information processing systems 30 (2017).

[40] Dahlgren, M. W. et al. Adventitial stromal cells define group 2 innate lymphoid cell tissue niches. Immunity 50, 707–722 (2019).

[41 ] Chen, T., Kornblith, S., Norouzi, M. & Hinton, G. A simple framework for contrastive learning of visual representations. Proceedings of Machine Learning Research 119, 1597–1607 (2020). URL https://proceedings.mlr.press/v119/chen20j.html.

[42] Pedregosa, F. et al. Scikit-learn: Machine learning in Python. Journal of Machine Learning Research 12, 2825–2830 (2011). URL https://jmlr.org/papers/v12/pedregosa11a.html.

[43] van der Walt, S. et al. scikit-image: image processing in Python. PeerJ 2, e453 (2014).

[44] Hu, W., et al. Strategies for pre-training graph neural networks. International Conference on Learning Representations (2020). URL https://openreview.net/forum?id=HJlWWJSFDH.

[45] Ba, J. L., Kiros, J. R. & Hinton, G. E. Layer normalization. *arXiv preprint arXiv:1607.06450* (2016).

[46] Xu, K., et al. Representation learning on graphs with jumping knowledge networks. Proceedings of Machine Learning Research 80, 5453–5462 (2018). URL https://proceedings.mlr.press/v80/xu18c.html.

[47] Liao, P.-S., Chen, T.-S. & Chung, P.-C. A fast algorithm for multilevel thresholding. Journal of Information Science and Engineering 17, 713–727 (2001).

[48] Sundararajan, M., Taly, A. & Yan, Q. Axiomatic attribution for deep networks. Proceedings of Machine Learning Research 70, 3319–3328 (2017).

