## Supplementary material for "Mantpy: a framework for extracellular matrix analysis in spatial proteomics": Combined supplementary

### Supplementary Information

**Supplementary Table 1 Related tools scored against the four capabilities required for spatial ECM analysis**

| Method | Input | Documented built-in capability | | | |
|---|---|---|---|---|---|
| | | (1) scverse | (2) ECM graph deep learning | (3) single- or multi-marker ECM graph | (4) cells and matrix in one graph |
| Matrisome AnalyzeR [1] | Bulk and single-cell omics | N | N | N | N |
| MatriCom [2] | Single-cell transcriptomics | N | N | N | N |
| MatriSpace [3] | Spatial transcriptomics | N | N | N | N |
| Pixie [4] | Multiplexed tissue images | N | N | N | N |
| Squidpy [5] | Spatial omics and tissue imaging | Y | N | N | N |
| GraphCompass [6] | Spatial proteomics and transcriptomics | Y | N | N | N |
| BANKSY [7] | Spatial omics (cells or spots) | Y | N | N | N |
| UTAG [8] | Multiplexed tissue images (cells) | Y | N | N | N |
| CellCharter [9] | Spatial omics (cells or spots) | Y | N | N | N |
| Bilgin *et al.* [10] | 3D collagen imaging | N | N | N | N |
| Bilgin *et al.* [11] | H&E histology | N | N | N | N |
| Grapa *et al.* [12] | Confocal fibronectin imaging | N | N | Y | N |
| **Mantpy** | Spatial proteomics | Y | Y | Y | Y |

**Supplementary Table 2 Mantpy modules**

| Module | Functionality | Key dependencies |
|---|---|---|
| Datasets `mt.datasets` | Three ECM-inclusive spatial proteomics datasets with loaders | — |
| Preprocessing `mt.pp` | ECM patch extraction, patch features and ECM clustering | Squidpy [5], SimCLR [13], scikit-learn [14], Leiden [15] |
| Graph building `mt.gr` | ECM, cell and cell–ECM graphs; grid, radius, $k$-NN, Delaunay and containing-patch rules per relation | Squidpy [5] |
| Analysis `mt.tl` | ECM co-occurrence, cell–ECM association and enrichment, graph statistics, label reconstruction | — |
| Graph learning `mt.nn` | PyTorch Geometric export; graph autoencoder and classifier; node importance | PyG [16], GraphMAE [17], GraphSAGE [18], integrated gradients [19] |
| Visualisation `mt.pl` | ECM and cell–ECM graph views, marker bubble plots, cell-focused ECM plots | — |

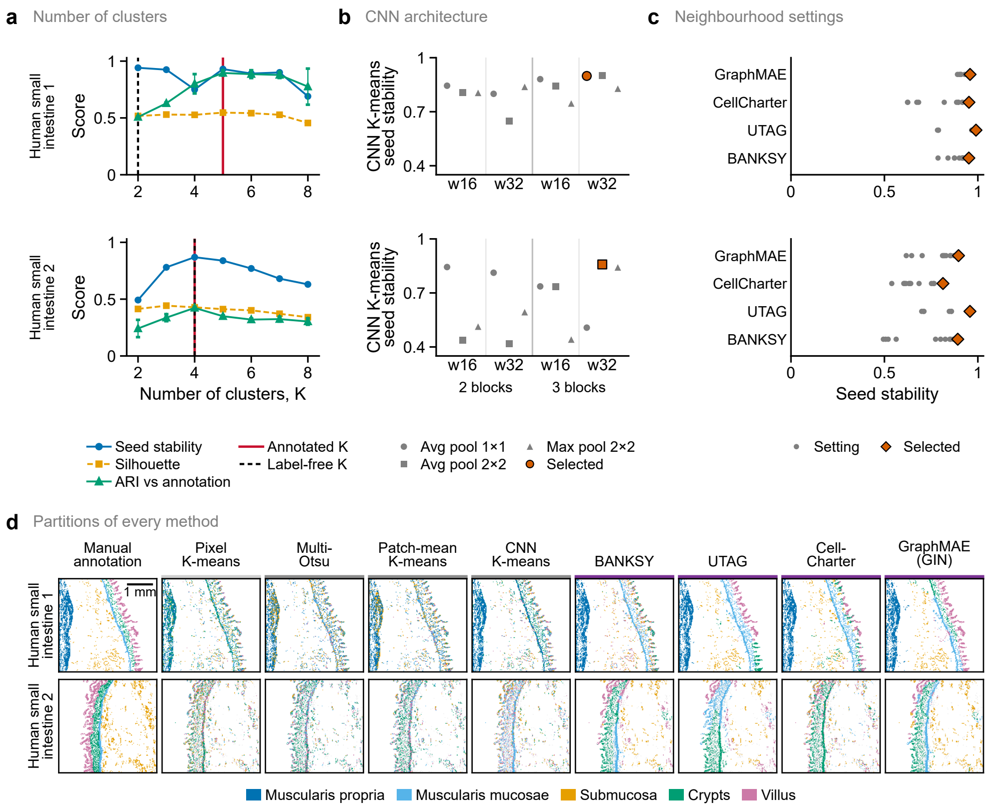


**Supplementary Fig. 1 Label-free choice of the number of clusters, CNN and neighbourhood settings, and partitions of every method in the two intestine images.** Rows, human small intestine 1 and 2. **a**, Number of clusters: seed stability (mean pairwise ARI of three training seeds), mean silhouette and ARI against the annotation (mean ± s.d. over seeds) at $K = 2$–8, on the GraphMAE embeddings of the selected configuration (Supplementary Table 3). Red line, annotated $K$, used in all analyses; dashed black line, label-free $K$ (highest stability). **b**, CNN architecture: CNN K-means seed stability of the twelve candidate CNNs (two or three convolutional blocks; width 16 or 32, w16 and w32; three readouts, marker shape). Vermilion, the most stable architecture, used in the tuned comparison (Fig. 2c, squares). **c**, Neighbourhood settings: seed stability of each spatial method's twelve settings on that CNN (grey points); vermilion diamonds, the most stable setting, used in Fig. 2c and **d**. GraphMAE uses a GIN encoder. Not shown: three UTAG settings per image that exceeded the edge-count guard and one image-2 UTAG setting whose archived labels are incomplete. **d**, Partitions of every method at the settings selected in **b**,**c** (seed closest to the three-seed median); intensity baselines, single runs. Clusters are matched one-to-one to annotation classes for display only. Light grey, annotated but not analysed (partitions) or analysed but not annotated (annotation); white, background. Bars above the maps: light grey, pixel; grey, patch; purple, graph methods. Scale bar, 1 mm. ARI, adjusted Rand index; CNN, convolutional neural network; GIN, graph isomorphism network; $K$, number of clusters.

**Supplementary Table 3 Label-free end-to-end search of the intestine GraphMAE pipeline**

| Component | Setting | Values searched | Selected | |
|---|---|---|---|---|
| | | | Image 1 | Image 2 |
| Graph construction | Patch size (px) | 16, 32, 64 | 32 | 32 |
| | $k$NN neighbours | 10, 20, 40, 80, 120, 160, 200, 240 | 120 | 160 |
| | Background-edge threshold | 0, 0.25, 0.5, 0.75 | 0.25 | 0.25 |
| CNN architecture | Convolutional blocks | 2, 3, 4 | 3 | 3 |
| | Width (channels) | 8, 16, 32 | 16 | 16 |
| | Readout | avg pool 2×2, avg pool 1×1, max pool 2×2, flatten | avg pool 2×2 | avg pool 2×2 |
| | Activation | ReLU, GELU | ReLU | ReLU |
| | Embedding dimension | 32, 64, 96, 128, 256 | 96 | 96 |
| | Projection head | 64, none | 64 | 64 |
| CNN training | NT-Xent temperature | 0.1, 0.2, 0.5 | 0.2 | 0.2 |
| | Augmentation | default, strong | default | default |
| | Learning rate | $3 \times 10^{-4}$, $10^{-3}$, $3 \times 10^{-3}$ | $10^{-3}$ | $10^{-3}$ |
| | Weight decay | $10^{-6}$, $10^{-5}$, $10^{-4}$ | $10^{-5}$ | $10^{-5}$ |
| | Epochs | 100, 200, 400 | 200 | 200 |
| | Batch size | 512, 1,024 | 1,024 | 1,024 |
| | Standardised embedding | yes, no | yes | yes |
| | Input window (× patch) | 1, 2 | 1 | 1 |
| Edge features | Distance encoding | log, raw, inverse, inverse square, Gaussian, none | log | log |
| | Orientation | yes, no | yes | yes |
| | Continuity | no, yes | no | no |
| | Encoder | GINE, GIN | GINE | GINE |
| GraphMAE | Layers | 1, 2, 3, 4, 5, 6 | 4 | 4 |
| | Width | 64, 128, 256 | 128 | 128 |
| | Mask ratio | 0.25, 0.5, 0.75 | 0.5 | 0.5 |
| | Learning rate | $3 \times 10^{-4}$, $10^{-3}$, $3 \times 10^{-3}$ | $10^{-3}$ | $10^{-3}$ |
| | Weight decay | $10^{-5}$, $10^{-4}$ | $10^{-4}$ | $10^{-4}$ |
| | Updates | 100, 200, 400 | 200 | 200 |
| | Standardised input | no, yes | no | no |

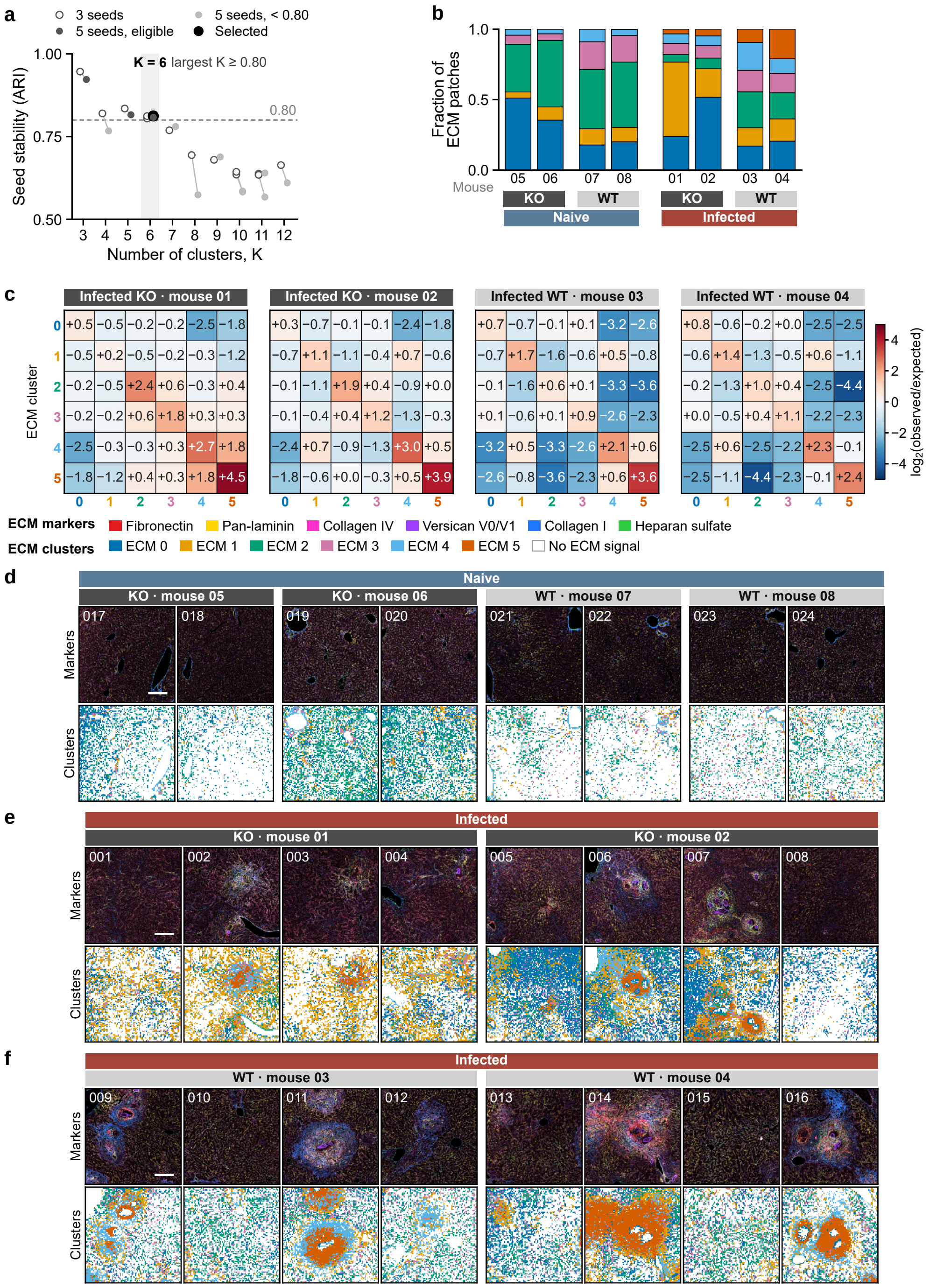


**Supplementary Fig. 2 Liver cluster selection, per-mouse composition and co-occurrence, and all 24 ROIs.** **a**, Number of clusters: seed stability (mean pairwise ARI) of the encoder–$K$ pairs within 0.01 of the best three-seed stability at each $K$ (open circles) after refitting with five seeds (filled; light grey, below 0.80); $K$ is the largest value whose five-seed stability reaches 0.80 ($K = 6$, 0.81; black). Search settings, Supplementary Table 4. **b**, Fraction of each mouse's ECM patches per cluster; Fig. 3a shows group means. **c**, $\log_2$(observed/expected) co-occurrence of each pair of ECM clusters in each infected mouse on the Delaunay graph of patch centres; Fig. 3c shows the KO − WT difference. **d**–**f**, All naive (**d**), infected KO (**e**) and infected WT (**f**) ROIs: six-marker composite (one display scale for all ROIs) and ECM clusters (white, no ECM signal); numbers, public ROI identifiers. Scale bars, 200 $\mu$m; each field is $1{,}000 \times 1{,}000$ pixels (1 pixel = 1 $\mu$m). Two mice per group; no tests. ARI, adjusted Rand index; KO, knockout; ROI, region of interest; WT, wild type.

**Supplementary Table 4 Label-free search of the liver encoder and number of clusters**

| Component | Setting | Values searched | Selected |
|---|---|---|---|
| Patches | Patch size (px) | 5, 10, 15, 20 | 10 |
| CNN architecture | Convolutional blocks | 2, 3, 4 | 2 |
| | Width (channels) | 8, 16, 32 | 32 |
| | Readout | avg pool 2×2, avg pool 1×1, max pool 2×2, flatten | max pool 2×2 |
| | Activation | ReLU, GELU | GELU |
| | Embedding dimension | 32, 64, 96, 128, 256 | 96 |
| | Projection head | 64, none | 64 |
| CNN training | NT-Xent temperature | 0.1, 0.2, 0.5 | 0.1 |
| | Augmentation | default, strong | default |
| | Learning rate | $3 \times 10^{-4}$, $10^{-3}$, $3 \times 10^{-3}$ | $3 \times 10^{-4}$ |
| | Weight decay | $10^{-6}$, $10^{-5}$, $10^{-4}$ | $10^{-4}$ |
| | Epochs | 100, 200, 400 | 200 |
| | Batch size | 512, 1,024 | 1,024 |
| | Standardised embedding | yes, no | no |
| Clustering | Number of clusters, $K$ | 3–12 | 6 |

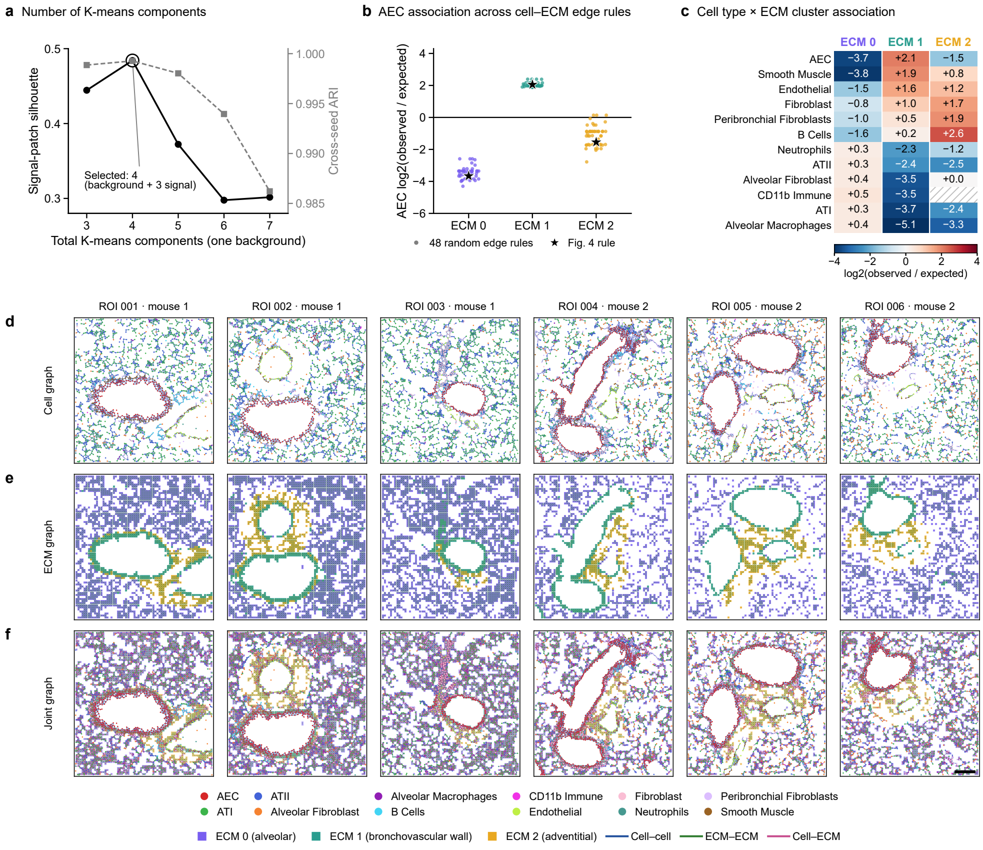


**Supplementary Fig. 3 ECM clusters, cell–ECM association and graphs of all six ROIs in healthy mouse lung.** Six regions of interest (ROIs) from two mice. **a**, Number of K-means components: signal-patch silhouette (black) and cross-seed ARI (grey) for three to seven components, one of them background; circle, the four selected. **b**, AEC $\log_2$(observed/expected) association with each ECM cluster, pooled over ROIs, under 48 random cell–ECM edge rules from the denoising search space (points) and the Fig. 4 rule (star). **c**, $\log_2$(observed/expected) association of every cell type with each ECM cluster, pooled over ROIs; colour clipped at ±4; hatched, no observed edges. **d**–**f**, Cell (**d**), ECM (**e**) and joint cell–ECM (**f**) graphs of all six ROIs, built with the rules of Fig. 4a–c. Scale bar, 100 $\mu$m. AEC, airway epithelial cell; ARI, adjusted Rand index; ATI and ATII, alveolar type I and II cells.

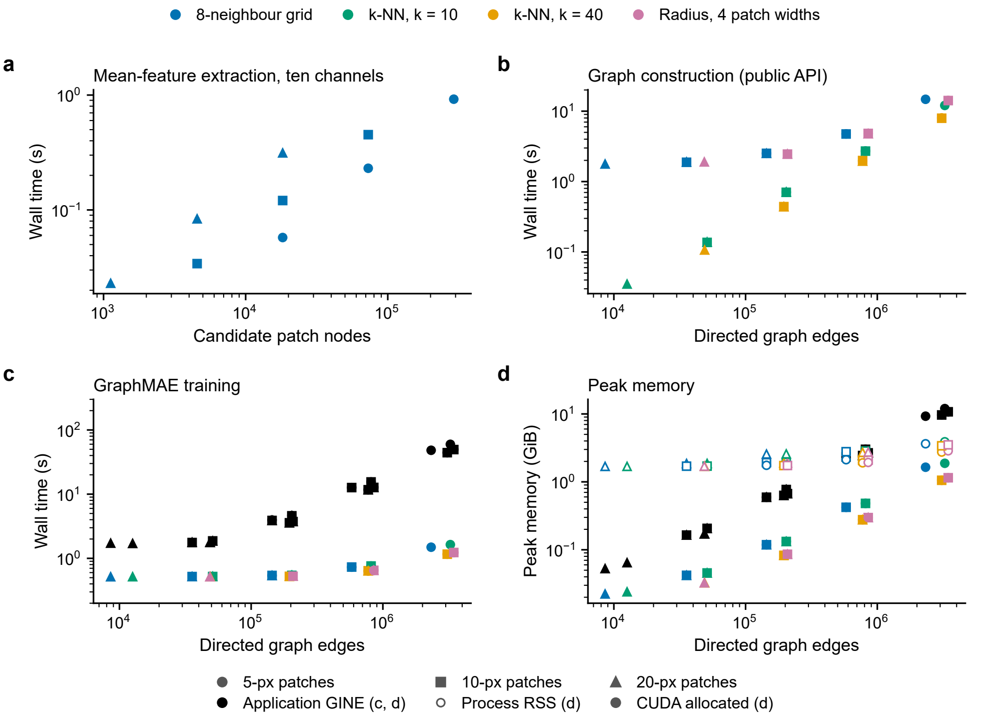


**Supplementary Fig. 4 Computational scaling of feature extraction, graph construction and GraphMAE training.** Profiles use one real $690 \times 660$-pixel ten-marker lung ROI and repeated $2 \times 2$ or $4 \times 4$ mosaics, 5-, 10- or 20-pixel patches and four graph rules, on an otherwise idle cloud GPU instance (NVIDIA A10G, 22 GiB; four CPU intra-op threads, one inter-op thread). All patches are retained and features are channel means. Points and error bars, mean $\pm$ sample s.d. of three serial fresh-process repeats. **a**, Feature-extraction time, measured on the grid graph only. **b**, Public graph-construction time versus directed edge count. **c**, GraphMAE training time: coloured, the public profile (two-layer, width-64 GIN without edge attributes, 20 updates, including setup); black, the application architecture of the intestine analysis (four-layer, width-128 GINE with three standardised edge features, 200 updates). **d**, Peak memory: sampled process RSS (open) and allocated CUDA memory (filled). The public profile completed 102 of 108 proposals; six exceeded the preallocation guard (4×4 mosaic, 5-pixel patches, $k$NN-40 and radius graphs; 12.3 and 13.9 million directed edges), on which the GINE model ran out of memory. Mosaics are computational stress tests, not biological validation; CNN fitting is excluded. RSS, resident set size.

**Supplementary Table 5 Nested-tuning configurations of the *Schistosoma*-liver classifier for each held-out mouse**

| Held-out mouse | Group | Graph | Layer type | Layers × width | Inner log-loss | $P$(infected) tuned | $P$(infected) untuned |
|---|---|---|---|---|---|---|---|
| 01 | Infected | radius 50 px | GraphSAGE | 4 × 256 | 0.301 | 0.935 | 0.823 |
| 02 | Infected | radius 40 px | GraphSAGE | 4 × 128 | 0.239 | 0.755 | 0.697 |
| 03 | Infected | radius 50 px | GraphSAGE | 4 × 128 | 0.263 | 0.832 | 0.634 |
| 04 | Infected | radius 40 px | GAT | 3 × 64 | 0.345 | 0.583 | 0.733 |
| 05 | Naive | radius 40 px | GraphSAGE | 4 × 128 | 0.299 | 0.248 | 0.422 |
| 06 | Naive | radius 50 px | GIN | 2 × 256 | 0.259 | 0.711 | 0.510 |
| 07 | Naive | radius 40 px | GraphSAGE | 4 × 128 | 0.251 | 0.405 | 0.447 |
| 08 | Naive | radius 40 px | GraphSAGE | 4 × 128 | 0.286 | 0.347 | 0.424 |

| Mouse | Training settings |
|---|---|
| 01 | mean aggregation; no normalisation; residual; GELU; dropout 0.25 (input 0.02); Adam, learning rate $2.1 \times 10^{-3}$, weight decay $3.2 \times 10^{-6}$; 300 epochs, constant schedule; label smoothing 0.02 |
| 02 | max aggregation; layer norm; residual; ReLU; dropout 0.01 (input 0.13); Adam, learning rate $2.8 \times 10^{-2}$, weight decay $1.4 \times 10^{-5}$; 500 epochs, constant schedule; label smoothing 0.04 |
| 03 | mean aggregation; no normalisation; no residual; ReLU; dropout 0.21 (input 0.22); Adam, learning rate $3.9 \times 10^{-3}$, weight decay $5.4 \times 10^{-6}$; 200 epochs, constant schedule; label smoothing 0.03 |
| 04 | 4 attention heads; batch norm; residual; GELU; dropout 0.27 (input 0.17); Adam, learning rate $2.0 \times 10^{-2}$, weight decay $2.1 \times 10^{-5}$; 300 epochs, cosine schedule; label smoothing 0.09 |
| 05 | mean aggregation; no normalisation; residual; ReLU; dropout 0.28 (input 0.14); Adam, learning rate $5.6 \times 10^{-3}$, weight decay $2.1 \times 10^{-5}$; 500 epochs, cosine schedule; label smoothing 0.03 |
| 06 | sum aggregation (GIN); batch norm; no residual; ReLU; dropout 0.35 (input 0.26); Adam, learning rate $4.7 \times 10^{-3}$, weight decay $1.7 \times 10^{-6}$; 200 epochs, cosine schedule; label smoothing 0.01 |
| 07 | mean aggregation; no normalisation; residual; GELU; dropout 0.05 (input 0.23); Adam, learning rate $8.4 \times 10^{-3}$, weight decay $1.6 \times 10^{-6}$; 300 epochs, constant schedule; label smoothing 0.02 |
| 08 | mean aggregation; no normalisation; residual; ReLU; dropout 0.02 (input 0.14); Adam, learning rate $1.5 \times 10^{-2}$, weight decay $4.7 \times 10^{-4}$; 500 epochs, constant schedule; label smoothing 0.01 |

**Supplementary Table 6 Nested-optimisation configurations of the label-only cell-context model for each held-out mouse and each held-out ROI**

| Held-out | Cell–cell | Cell–ECM | Edge weight | Composition | Penalty ($C$) | Cleaning rule | Calibration / inner accuracy |
|---|---|---|---|---|---|---|---|
| *Mouse held out (main result; inner loop over the training mouse's three ROIs, 60 trials)* | | | | | | | |
| Mouse 1 | radius 15 px | radius 15 px | Gaussian | count | elasticnet (1.11) | $-\log P > 2.99$; confidence $\geq 0.5$; gate none | none / 0.951 |
| Mouse 2 | Delaunay | 9-NN, ≤20 px | inverse distance | fraction | l1 (31.067) | $1 - P > 0.95$; confidence $\geq 0.0$; gate none | none / 0.967 |
| *ROI held out (inner loop over the five training ROIs, 150 trials)* | | | | | | | |
| ROI 001 | 10-NN, ≤40 px | Delaunay | Gaussian | log(1+count) | l1 (0.034) | $1 - P > 0.93$; confidence $\geq 0.5$; gate none | temperature / 0.963 |
| ROI 002 | radius 39 px | containing patch + 8 neighbours | Gaussian | log(1+count) | elasticnet (0.774) | margin $> 0.86$; confidence $\geq 0.7$; gate none | none / 0.965 |
| ROI 003 | Delaunay | containing patch + 8 neighbours | none | log(1+count) | l2 (0.01) | $-\log P > 2.92$; confidence $\geq 0.5$; gate none | temperature / 0.959 |
| ROI 004 | Delaunay | 8-NN | none | log(1+count) | l1 (10.242) | min–max $-\log P > 0.28$; confidence $\geq 0.7$; gate none | temperature / 0.966 |
| ROI 005 | Delaunay | containing patch + 8 neighbours | none | fraction | l2 (0.16) | $1 - P > 0.94$; confidence $\geq 0.7$; gate none | temperature / 0.967 |
| ROI 006 | Delaunay | Delaunay, ≤60 px | inverse distance | fraction | elasticnet (34.338) | $1 - P > 0.95$; confidence $\geq 0.0$; gate none; smoothing | temperature / 0.956 |

### References


[1] Petrov, P. B., Considine, J. M., Izzi, V. & Naba, A. Matrisome AnalyzeR–a suite of tools to annotate and quantify ECM molecules in big datasets across organisms. *Journal of Cell Science* **136**, jcs261255 (2023).

[2] Lamba, R., Paguntalan, A. M., Petrov, P. B., Naba, A. & Izzi, V. MatriCom, a single-cell RNA-sequencing data mining tool to infer cell–extracellular matrix interactions. *Journal of Cell Science* **138**, jcs263927 (2025).

[3] Oshinjo, A., Chen, D., Petrov, P., Izzi, V. & Naba, A. MatriSpace: Identification and visualization of spatially resolved ECM gene expression patterns in health and disease. *bioRxiv* (2026). URL https://doi.org/10.64898/2026.04.26.720198. Preprint; not peer reviewed.

[4] Liu, C. C. *et al.* Robust phenotyping of highly multiplexed tissue imaging data using pixel-level clustering. *Nature Communications* **14**, 4618 (2023).

[5] Palla, G. *et al.* Squidpy: a scalable framework for spatial omics analysis. *Nature Methods* **19**, 171–178 (2022).

[6] Ali, M. *et al.* GraphCompass: spatial metrics for differential analyses of cell organization across conditions. *Bioinformatics* **40**, i548–i557 (2024).

[7] Singhal, V. *et al.* BANKSY unifies cell typing and tissue domain segmentation for scalable spatial omics data analysis. *Nature Genetics* **56**, 431–441 (2024).

[8] Kim, J. *et al.* Unsupervised discovery of tissue architecture in multiplexed imaging. *Nature Methods* **19**, 1653–1661 (2022).

[9] Varrone, M., Tavernari, D., Santamaria-Martínez, A., Walsh, L. A. & Ciriello, G. CellCharter reveals spatial cell niches associated with tissue remodeling and cell plasticity. *Nature Genetics* **56**, 74–84 (2024).

[10] Bilgin, C. C., Lund, A. W., Can, A., Plopper, G. E. & Yener, B. Quantification of three-dimensional cell-mediated collagen remodeling using graph theory. *PLOS ONE* **5**, e12783 (2010).

[11] Bilgin, C. C., Bullough, P., Plopper, G. E. & Yener, B. ECM-aware cell-graph mining for bone tissue modeling and classification. *Data Mining and Knowledge Discovery* **20**, 416–438 (2010).

[12] Grapa, A.-I., Efthymiou, G., Van Obberghen-Schilling, E., Blanc-Féraud, L. & Descombes, X. A spatial statistical framework for the parametric study of fiber networks: Application to fibronectin deposition by normal and activated fibroblasts. *Biological Imaging* **3**, e25 (2023).

[13] Chen, T., Kornblith, S., Norouzi, M. & Hinton, G. A simple framework for contrastive learning of visual representations. *Proceedings of Machine Learning Research* **119**, 1597–1607 (2020). URL https://proceedings.mlr.press/v119/chen20j.html.

[14] Pedregosa, F. *et al.* Scikit-learn: Machine learning in Python. *Journal of Machine Learning Research* **12**, 2825–2830 (2011). URL https://jmlr.org/papers/v12/pedregosa11a.html.

[15] Traag, V. A., Waltman, L. & van Eck, N. J. From Louvain to Leiden: guaranteeing well-connected communities. *Scientific Reports* **9**, 5233 (2019).

[16] Fey, M. & Lenssen, J. E. Fast graph representation learning with PyTorch Geometric. *arXiv preprint arXiv:1903.02428* (2019).

[17] Hou, Z. *et al.* GraphMAE: Self-supervised masked graph autoencoders. *Proceedings of the 28th ACM SIGKDD Conference on Knowledge Discovery and Data Mining* 594–604 (2022).

[18] Hamilton, W., Ying, Z. & Leskovec, J. Inductive representation learning on large graphs. *Advances in neural information processing systems* **30** (2017).

[19] Sundararajan, M., Taly, A. & Yan, Q. Axiomatic attribution for deep networks. *Proceedings of Machine Learning Research* **70**, 3319–3328 (2017).
